# Comprehensive integration of single-cell transcriptomic data illuminates the regulatory network architecture of plant cell fate specification

**DOI:** 10.1101/2022.10.24.513543

**Authors:** Shanni Cao, Chao He, Xue Zhao, Ranran Yu, Yuqi Li, Wen Fang, Chen-Yu Zhang, Wenhao Yan, Dijun Chen

**Affiliations:** State Key Laboratory of Pharmaceutical Biotechnology, School of Life Sciences, Nanjing University, Nanjing 210023, China; National Key Laboratory of Crop Genetic Improvement, Hubei Hongshan Laboratory, Huazhong Agricultural University, Wuhan, China

## Abstract

Plant morphogenesis relies on precise gene expression programs at the proper time and position which is orchestrated by transcription factors (TFs) in intricate regulatory networks at a cell-type specific manner. Here we presented a reference single-cell transcriptomic atlas of *Arabidopsis* seedlings by integration of 40 published scRNA-seq datasets from representative tissues as well as the entire under- and above-ground parts. We identified 34 distinct cell types or states, largely expanding our current view of plant cell compositions. We then mapped the developmental trajectory of root-shoot lineage separation and identified differential gene expression programs that may regulate the cell fate determination of under- and above-ground organs. Lastly, we systematically constructed cell-type specific gene regulatory networks and uncovered key regulators that act in a coordination manner to control cell-type specific gene expression. Taken together, our study not only offers a valuable resource plant cell atlas exploration but also provides molecular insights into gene-regulatory programs that determines organ specify, particularly the differentiation between root and shoot.

## Introduction

Plant development starts with the embryogenesis followed by postembryonic development of above-ground tissues and roots of the plant body from the shoot and root apical meristems (SAM and RAM), respectively (Kaufmann et al., 2010; Wolters and Jürgens, 2009). Primary meristems (including shoot and root meristems) are established during the embryogenesis while the formation of secondary meristems (such as axillary meristems, the cambium and inflorescence meristems) is modulated during post-embryonic development (Wang et al., 2016; Melzer et al., 2008). The activity of all kinds of meristems enables plants to produce new organs and structures throughout their ontogeny, thus sustaining plant growth and architecture (Heidstra and Sabatini, 2014). Plant organ development involves continuing cell expansion and dynamics of cell differentiation with new functions, which relies on precise gene expression programs at the proper time and position. Plant organ is a mixture of multiple cell types with distinct features and fate, which are orchestrated by transcription factors (TFs) in intricate regulatory networks at a cell-type specific manner. Triggered by internal and external signals, TFs bind to *cis*-elements of downstream genes to regulate gene expression to further fulfil cell identity establishment and maintenance. Gene regulatory networks (GRNs) containing TFs and their target genes shed light on how genes co-ordinately orchestrate cell fate specification. Therefore, network-based approaches can help elucidate the mechanisms that link genes, cells, tissues and organs in a systemic manner(Wu et al., 2022).

GRN analyses that integrate multiple data types at increased resolution have significantly improved our understanding of the complex molecular mechanisms controlling the development of flowers (Chen et al., 2018; Yan et al., 2016), roots (Reynoso et al., 2022; Brady et al., 2011; Moreno-Risueno et al., 2015), seeds (Santos-Mendoza et al., 2008), and secondary cell walls (Taylor-Teeples et al., 2015). The rapid development of single cell sequencing techniques which have been fully embraced by the plant community (Seyfferth et al., 2021), has greatly promoted in-deep knowledge for plant organ morphogenesis at single-cell resolution. In fact, GRNs constructed at single-cell level revealed novel mechanisms in the root morphogenesis process in *Arabidopsis* (Yang et al., 2021; Denyer et al., 2019; Roszak et al., 2021). This suggests that sing-cell GRN (scGRN) analysis can recapitulate the complex and heterogeneous biological processes in the plant (Qian and Huang, 2020; Tripathi and Wilkins, 2021). Although progress has been made on constructing cell transcriptomic atlases in plants(Seyfferth et al., 2021), our current view of cellular taxonomy is restricted in specific organs or tissues, hampering our holistic understanding of plant cell fate specification. In fact, tremendous efforts from human studies have demonstrated that integration multiple single-cell transcriptomic datasets can boost the statistical power for discovery of rare cell phenotypes and novel markers (Stuart et al., 2019), highlighting the potential value of large-scale single-cell data integration.

In the current study, we presented for the first time a reference single-cell transcriptomic atlas of the whole *Arabidopsis* seedlings by integration of 40 single cell transcriptomic datasets from representative tissues. In total, 34 distinct cell types have been identified from 155,889 cells, expanding our current view of plant cell taxonomy. We defined a cell marker gallery which contains numbers of potentially new markers. Furthermore, we mapped the bifurcation of root-shoot developmental trajectory, explored the differential expressed genes that could be vital for cell fate determination of organs from different parts of the plant body. Lastly, we systematically constructed cell-type specific GRNs and uncovered key regulators that act in a coordination manner to control cell-type specific gene expression. Therefore, our integrative analysis will not only be a valuable resource for future plant single cell atlas construction, but also provide new insights for developmental regulatory mechanism at specific cell types.

## Results

### Comprehensive integration of single-cell transcriptomic data in Arabidopsis juvenile seedlings

To generate a reference single-cell transcriptomic atlas of the whole *Arabidopsis* seedling and thus to reconstruct the cellular taxonomy across the plant body, we performed comprehensive integration analysis of single-cell RNA sequencing (scRNA-seq) data generated from representative tissues in *Arabidopsis* juvenile seedlings (**Figure 1A-B**; see **Methods**). A total of 40 scRNA-seq datasets were collected from ten different studies (Liu et al., 2020; Lopez-Anido et al., 2021; Denyer et al., 2019; Wendrich et al., 2020; Zhang et al., 2019; Long et al., 2021; Jean-Baptiste et al., 2019; Ryu et al., 2019; Shulse et al., 2019; Farmer et al., 2021) (**Supplemental Data Set S1**). These data were generated from root tips, whole root, aerial tissues and cotyledons of *Arabidopsis* seedling plants at 5-10 days after germination (DAG), representing both under- and above-ground part of the whole plant (**Figure 1A**). After stringent quality control and removal of low-quality cells and potential doublets, a total of 155,889 high-quality cells were retained for further analysis (**Supplemental Data Set S2**). Unsupervised clustering based on uniform manifold approximation and projection (UMAP) reveals that shoot and root tissues have common and unique cell populations (**Figure 1B**). Overall, 46 distinct cell clusters were identified in the integrated cell map (**Figure 1C**). The number of cells in each cluster ranges from 220 (cluster 45, C45) to 10,007 (C0). It turns out that all the identified cell clusters are supported by different datasets from at least two different studies (**Figure 1D**), suggesting the high confidence of cell cluster identification. We further assigned the 46 clusters into three different groups based on the proportion of cells belonging to either the root- or shoot-related tissues. Using a threshold of 25%, we defined 11 shoot-specific clusters, 20 root-specific clusters and 15 shoot-root shared clusters (**Figure 1D**). We performed differential gene expression analysis among the three groups and identified highly expressed genes in each group of cell clusters (**Figure 1E** and **Supplemental Data Set S3**). Photosynthetic-related, salicylic and jasmonic acid signalling related pathways are enriched in shoot-specific clusters; while root-specific clusters showed high enrichment for biological pathways related to nutrition absorption and transport. In contrast, the shoot-root shared clusters are enriched for basic cellular processes such as RNA modification and ATP biosynthesis (**Figure 1E**). Taken together, the above data provide a full-view of cellular landscape of the whole *Arabidopsis* plant at the juvenile stage.

**Figure 1:**
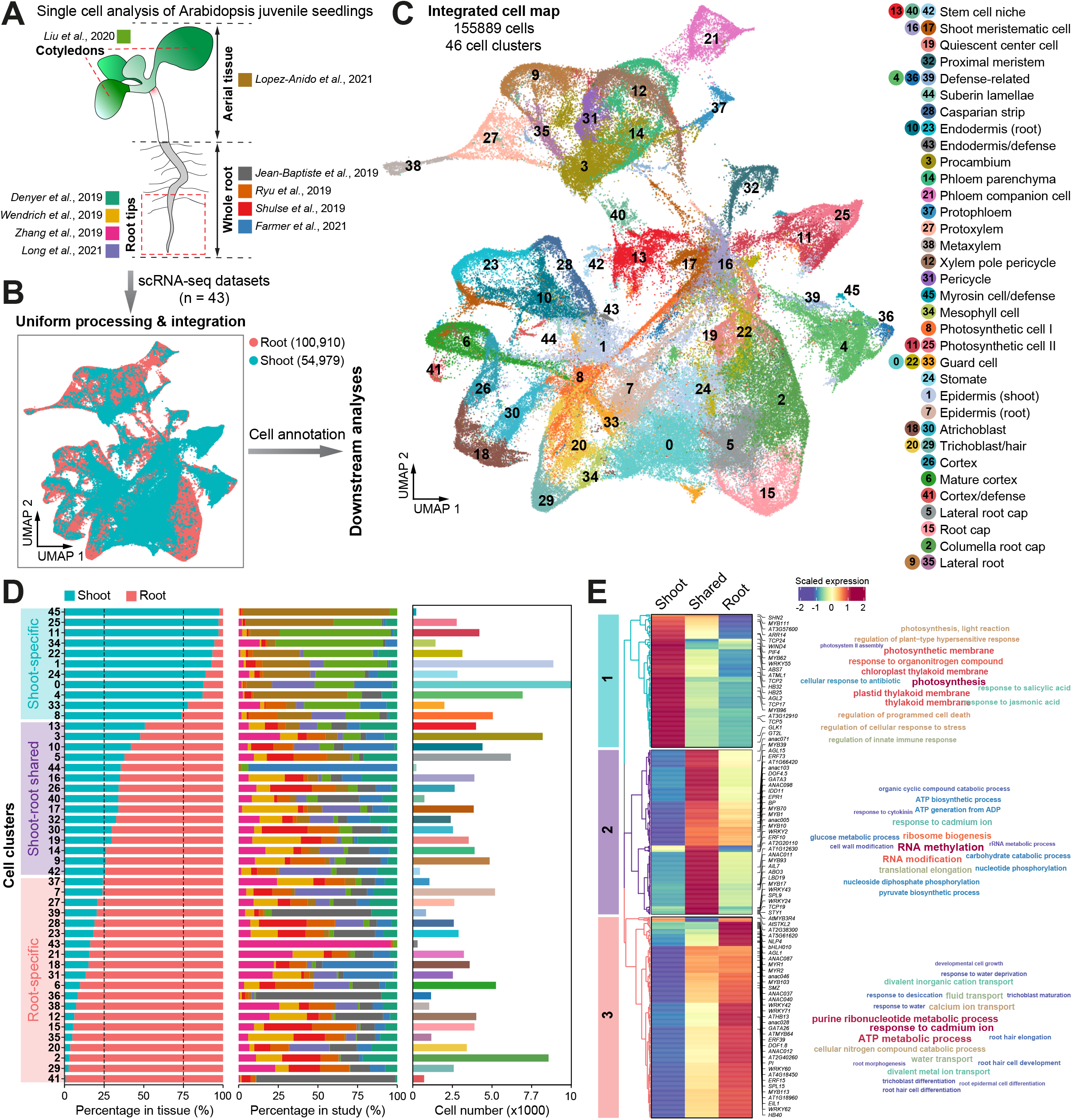
Mapping a reference single-cell transcriptomic atlas of *Arabidopsis* seedlings at the whole-organism level. **(A)** Single cell transcriptomic data used for constructing the reference single-cell transcriptomic atlas. In total, 40 scRNA-seq datasets were collected from ten different studies (in different color codes). The datasets were either generated in shoot or root tissues, as annotated according to the original studies. **(B)** Bioinformatic analysis pipeline for integrative analysis of single cell datasets. All scRNA-seq datasets were subjected to uniform processing, quality control and integration. The Uniform Manifold Approximation and Projection (UMAP) plot shows the integrated cell map where cells are colored by the tissues of origin. **(C)** UMAP plot displaying the integrated cell map, which consists of 46 distinct cell clusters. Cell clusters were annotated based on canonical marker genes, as shown in **Figure 2A**. **(D)** Bar plots displaying the distribution of cells in each cluster based on the tissue of origin (left) or the original study (middle; color codes as in **A**), and the number of cells (right; color codes as in **C**). Cell clusters are assigned to three different categories based on the composition of cells from shoot or root tissues. **(E)** Heatmap showing the highly expressed genes in the three different categories of cell clusters (left). Representative genes are highlighted on the right. Word cloud graphs (right) indicate enriched gene ontology (GO) biological progresses for genes in the three categories.

### Systemic annotation of Arabidopsis cell types from integrated datasets

To test the ability of the integrated single-cell transcriptomic map for identifying rare and novel cell types, we dedicated to annotate each cell cluster based on their top differentially expressed genes (DEGs; **Supplemental Data Set S3**). In general, these DEGs are highly cluster specific (**Figure 2A-B**) and are thus valuable for cell type annotation. We carefully determined the cell type of each cluster using known markers collected from existing single-cell studies (Jean-Baptiste et al., 2019; Wendrich et al., 2020; Long et al., 2021; Farmer et al., 2021; Shahan et al., 2022; Lopez-Anido et al., 2021; Shulse et al., 2019; Zhang et al., 2019; Denyer et al., 2019; Liu et al., 2020; Zhang et al., 2021) or information of cell identity provided by fluorescence in situ hybridization (FISH) from literature (**Supplemental Figure S1** and **Supplemental Data Set S4**). In total 34 distinct cell types were identified and they can be grouped into nine different major types (cortex, endodermis, epidermis, defence-related cells, initiation cells, photosynthetic cells, root cap, stele cells and stomata cells) according to their functions or spatial locations on the UMAP (**Figures 1C** and **2A**; **Supplemental Note**). To the best of our knowledge, our analysis here provides the most comprehensive cell type annotation so far at single cell resolution, largely expanding our current view of cell compositions in *Arabidopsis* (Shahan et al., 2022; Long et al., 2021; Farmer et al., 2021; Ryu et al., 2019; Zhang et al., 2019). In addition, we are able to identify an extensive list of new marker genes based on the integrated cell atlas (**Figure 2A-B** and **Supplemental Data Set S4**). Notably, several novel cell types were identified in our integrative analysis, including suberin lamellae (Cluster 44), defense-related cells (C4, C36 and C39) and myrosin cells (C45) (**Figure 1C**). In *Arabidopsis* roots, suberin lamellae and casparin strips are both located in the endodermis and function as transmembrane barriers to limit the movement of apoplastic solutes into the endodermal cells (Franke and Schreiber, 2007). However, there are currently no available markers to distinguish these two cell types. We observed that *MYB39* (*AT4G17785*), *ABCG2* (*AT2G37360*) and *CYP86A1* (*AT5G58860*), which are involved in suberization of root endodermis(Cohen et al., 2020) and suberin monomer biosynthesis (Yadav et al., 2014; Vishwanath et al., 2015), were specifically expressed in C44 (**Figure 2B**). Therefore, the rare cell cluster C44 was annotated as suberin lamellae cells. Meanwhile, we identified C28 as casparin strip cells since its marker genes *CASP1-5* and *PER64* (**Figure 2B**) are essential for casparin strips formation (Lee et al., 2013). Interestingly, the majority of genes that explicitly expressed in the cluster of C4, C36 and C39 are heat shock proteins (HSP) and heat shock factors (HSF) transcription factors (TFs) (**Figure 2A**). Accordingly, functional enrichment analysis implies that the highly expressed genes from these cell clusters involved in the maintenance of cellular homeostasis and response to various environmental stimulus (**Supplemental Figure S2**). Therefore, we defined these three cell clusters as putative defense-related cells, whose biological function requires further experimental validation.

**Figure 2:**
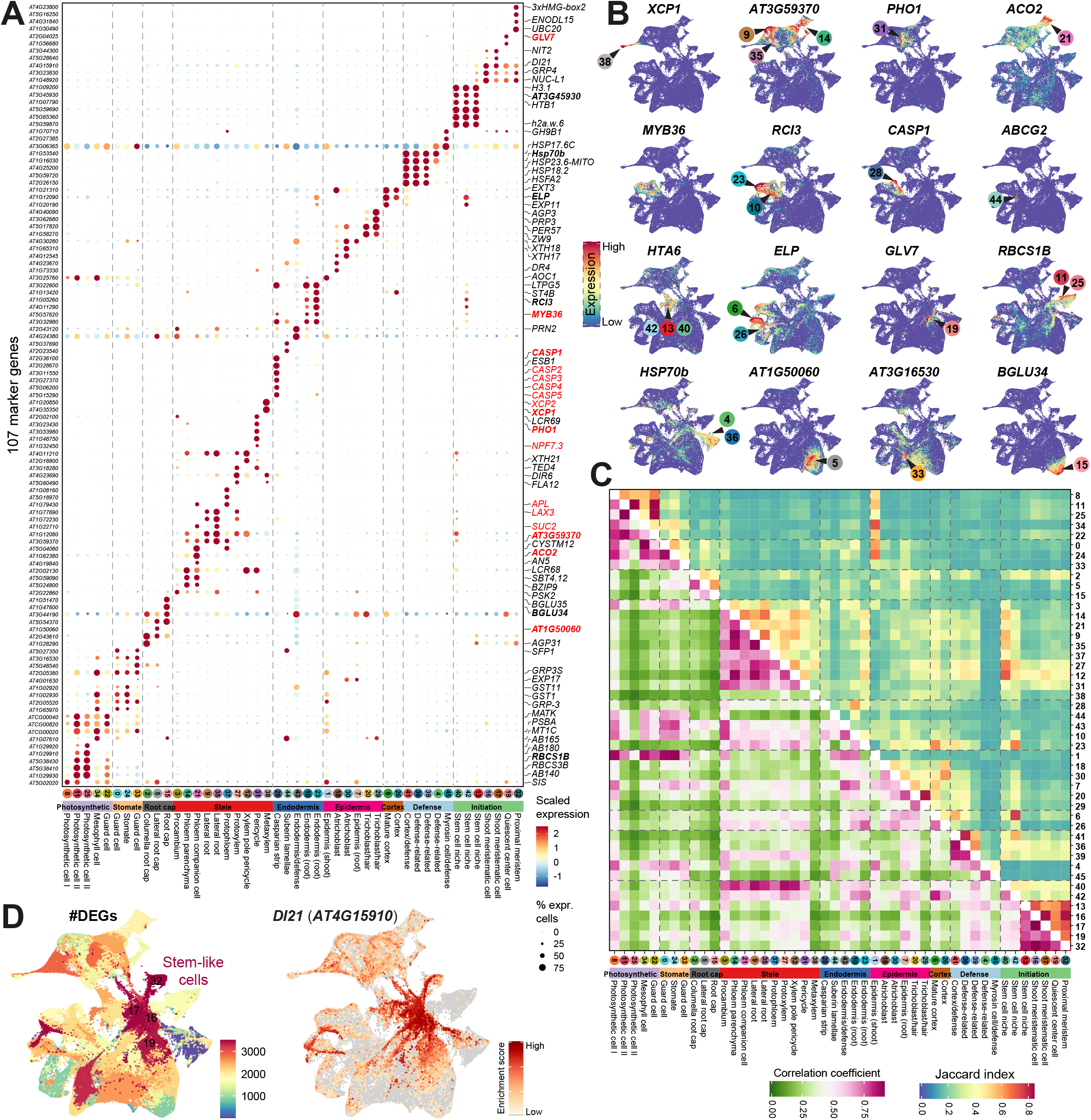
Comprehensive annotation of cell types and identification of new marker genes. **(A)** Dot plots showing the expression patterns of top marker genes (row) in each cell cluster (column). Cell types or states were annotated based on marker genes and can be roughly assigned to nine different major cell classes according to their structure and/or function. Gene accessions are shown on the left and representative markers are labeled on the right. Genes in red are candidates of new markers. Genes in bold are examples in (**B**). The dot size is proportional to the fraction of cells which express specific genes. The color scale corresponds to the relative expression of specific genes. (**B**)UMAP plots demonstrating the expression patterns of examples of marker genes. The corresponding cell cluster(s) where the marker gene was highly expressed are highlighted. (**C**)Heatmap showing the transcriptome similarity of cell clusters. Cell clusters are organized in the same order in (**A**). (**D**)UMAP plots displaying the number of differentially expressed genes (DEGs) among different cell clusters (left) or the expression pattern of *DI21* (*AT4G15910*; right).

To validate the confidence of cell type annotation, we used CIBERSORTx (Newman et al., 2019) to deconvolve cell-type specific gene expression from published bulk RNA-seq data that were generated from various tissues, organs or cell types in *Arabidopsis* seedlings (**Supplemental Data Set S5**). Generally, the estimated cell-type specific gene expression patterns from bulk data are consistent with the samples of origin (**Supplemental Figure S3**). For instance, photosynthetic cells and the corresponding marker genes were overrepresented in aboveground samples. Samples enriched for quiescent centre and stem cell niche showed dominant abundance of initiation cell clusters. The above analyses further confirm the reliability of cell-type annotation based on marker genes.

Furthermore, we evaluated the transcriptome similarity of different cell clusters by measuring the Jaccard similarity of DEGs between any two clusters as well as the overall expression similarity among clusters (**Figure 2C**). In general, cell clusters in the same major types showed significantly higher similarity than clusters from different major types (**Supplemental Figure S4**), suggesting that cells from different major types harbour distinct active gene expression programs. Indeed, the enrichment of biological processes for each cluster based on their DEGs were distributed into representative functional topics in a major type manner using the topic modelling program *CellFunTopic* (see **Methods**; **Supplemental Figure S5A**). For instance, chlorophyll biosynthetic process and response to light stimulus were highly enriched in the photosynthetic cell clusters (including C8, C11, C22 and C25) (**Supplemental Figure S5B**). Of note, we observed that four of the initiation cell clusters (C16, C17, C19 and C32) are significantly distinct from other clusters in terms of transcriptome similarity (**Figure 2C**). Accordingly, these clusters have the greatest number of DEGs (**Figure 2D**). Interestingly, these clusters coincidentally shared in both shoot- and root-related tissues (**Figure 1D**). Furthermore, we found that genes with roles in regulating meristem cell fate, such as *BBM* (PLT4) (Burkart et al., 2022), *PLT1* (Sarkar et al., 2007) and *DI21, were* highly expressed in these cell clusters (**Figure 1D** and **Supplemental Figure S6**). The above results indicate that these four initiation cell clusters may form the primary meristems in *Arabidopsis* (Stahl and Simon, 2010).

### Dynamic changes of gene expression programs underpinning the ontogeny of root and shoot tissues

In *Arabidopsis*, the developmental trajectories of root or shoot tissue lineages have been reconstructed independently in recent studies (Zhang et al., 2019; Lopez-Anido et al., 2021; Denyer et al., 2019). However, little is known about how differentiation orientation is controlled in space and time to produce and maintain the above- and belowground tissues. To decipher the developmental trajectory from the primary meristem cells to a diverse set of cell types from plant tissues (**Figure 3A**), we performed pseudotime trajectory analysis using all types of cells, which were arranged into a trajectory with three main bifurcations and five cell states (**Figure 3B**). Initiation cells including the QC (C19) and meristem cells (C16, C17 and C32) were located in the centre of the developmental trajectories (state 1). One bifurcation of the trajectories led to the formation of aboveground cells (state 2) including apical epidermis, guard cells (C0, C22 and C33), stomata (C24), photosynthetic cells (C8, C11 and C25) and mesophyll cells (C34). The second bifurcation led to the construction of root morphogenesis (CS, trichoblast, atrichoblast and lateral root) and some root-shoot shared cells including cortex, epidermis and vascular cells (state 3). The last one was responsible for the belowground part including root cap cells (state 4). Interestingly, state 5, which is close to state 1 and on the branch of trajectory of state 2, mainly contain the defence-related cells (C4) and photosynthetic cells (**Figure 3B-C**). Differentially expressed genes varying along pseudotime across the five states and their GO enrichment analysis suggest that transition from initiation cells to underground and aboveground cells are two distinct biological process (**Supplemental Figure S7**). In line with the trajectory inference, GO enrichment analysis suggests that genes highly expressed in state 2 and state 5 were associated with the photosynthesis associated pathways, while genes in state 1 were enriched in cell cycle associated pathways which echoes the role of initiation cells. State 3 was enriched with genes involved in the process of secondary cell wall formation and transport, associated to the differentiation of casparin strips, lateral root and root-shoot shared vascular cells. Genes engaged in the formation of root cap cells (state 4) were mostly nucleotide-sugar transmembrane transporter (**Figure 3D**).

**Figure 3:**
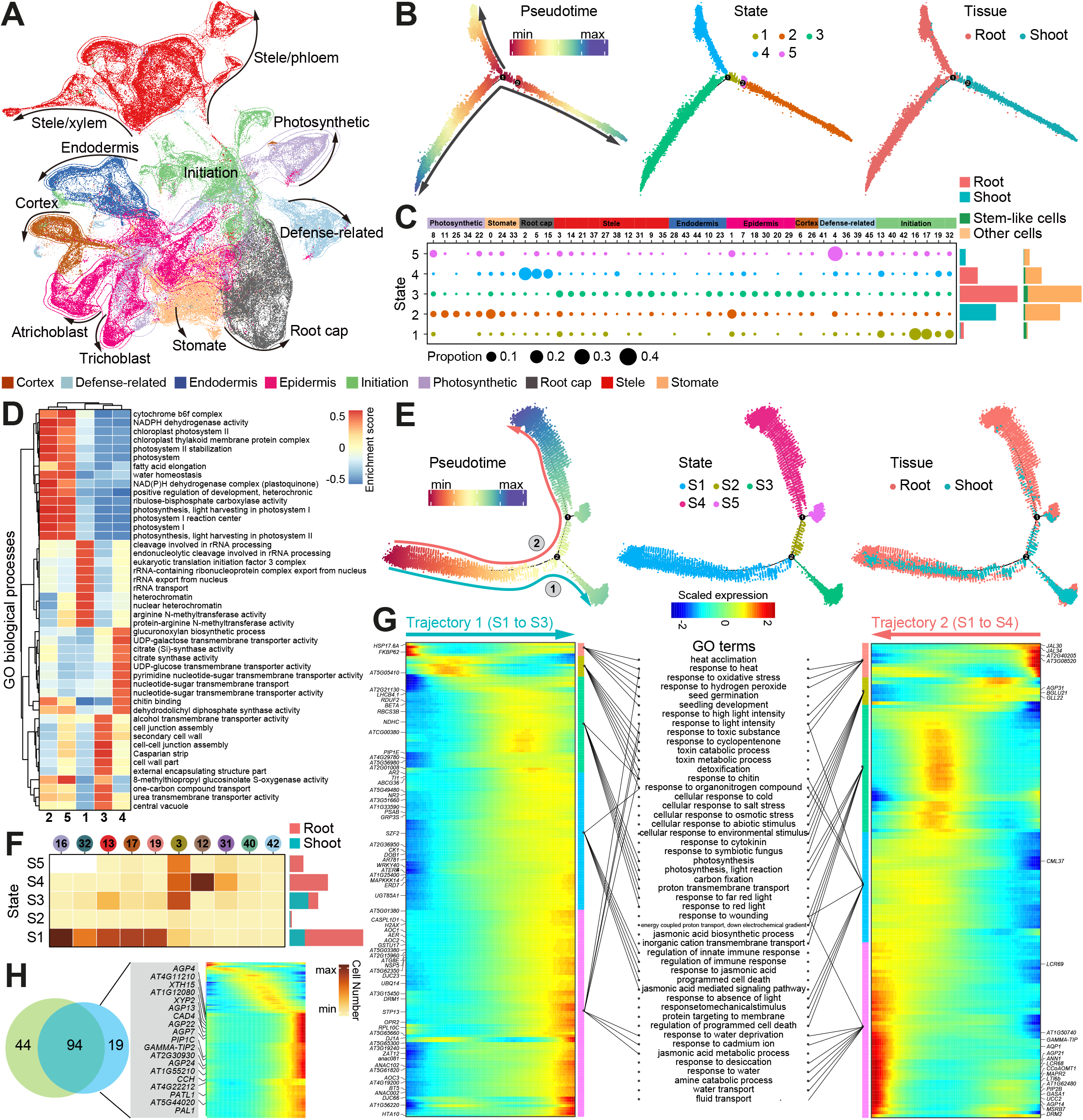
Semisupervised pseudotime analysis revealing of distinct developmental trajectories and associated gene programs underling root-shoot separation. (**A**)UMAP plot revealing overall organization similarity of major cell classes on the cell map. (**B**)Monocle2 pseudotime analysis of all major cell classes depicting distinct differential trajectories of initiation cells toward shoot and root-specific cells. Cells are ordered in pseudotime along the trajectory (left) and can be grouped into five different cell states (middle); cells with tissue origins are projected on the trajectory (right). (**C**)Bubble plot showing the proportion of different types of cells in each cell state. The right bars the number of cells in each state. (**D**)Heatmap displaying the enrichment of GO biological processes across cell states in (**B**). (**E**)Monocle2 pseudotime analysis of initiation cells revealing two branches of trajectories. The trajectory is colored based on pseudotime (left), cell state (middle) or tissue of origin (right). (**F**)Heatmap showing the scaled expression of genes that were differentially expressed across the two branches of pseudotime trajectories in (**E**). In each trajectory, genes are grouped in five distinct clusters according their expression patterns. The top three enriched biological pathways each cluster are shown in the middle panel. Shared and specific biological pathways for each trajectory are indicated. (**H**) Venn diagram showing the genes contributed to the development of root initiation cells that were shared between the developmental trajectories of root pericycle and procambium.

The above analysis reveals that the above- and belowground cells are differentiated from a pool of initiation cells through different trajectories. We further explored how cell fates are determined in the initiation cells using pseudotime trajectory analysis. Since pericycle and procambium cells have been shown to harbour meristematic features (Sugimoto et al., 2010; Atta et al., 2009; Parizot et al., 2008), they were also included in the analysis. We set the primary meristem cells (C16, C17, C19 and C32) as the start of the inferred trajectory. Two major branches of cells were found to derive from a common ancestral cell state, which consists of cells equally from both root and shoot tissues (**Figure 3E**), suggesting that the root and shoot meristems may share similar gene expression programs for cell differentiation. One major branch (trajectory 1) led to the differentiation of shoot procambium cells from the primary meristem cells while the other branch (trajectory 2) differentiating toward root procambium/pericycle cells (**Figure 3E-F**). Note that the trajectory 2 has two minor paths with different cell states: one was the root xylem pole pericycle (state S4) and the other still maintained as procambium (state S5) (**Figure 3E-F**).

We sought to examine gene expression programs driving the developmental trajectories of initiation cells by performing differential gene expression analysis over pseudotime (**Figure 3G**). During the development of shoot procambium (trajectory 1) and root procambium/pericycle cells (trajectory 2), genes involved in indole acetic acid (IAA) signalling pathway were activated, including *ABCG36, AIR12, BT5* and *XTH22* (**Figure 3G**). The result supports the notion that the IAA signal pathway is triggered to regulate the development of procambium/pericycle cells in both root and shoot (Zhang et al., 2022; Smetana et al., 2019; Vanneste et al., 2005). GO enrichment analysis showed that stress-response related GO terms are highly enriched in DEGs over the two trajectories but the enriched biological pathways are generally different between trajectories 1 and 2. In the shoot procambium development (trajectory 1), toxic and detoxification stress-response pathways were overrepresented, while in the root procambium developing process, cellular response to salt, osmotic and fungus were more dominant (**Figure 3G**). The observation is consistent with the totally different environment where shoot and root tissues are developing. As expected, photosynthesis and light response genes (*LHCB4*.*1* and *RBCS3B*) were found in the shoot procambium developing process, while genes related to root morphogenesis (*AGP14* and *CML37*), water uptake (*PIP2B* and *GAMMA-TIP*) and root epidermal cell differentiation (*ANN1*) were found in the root procambium development (**Figure 3G**). In addition, most of the genes contributed to the development of root initiation cells were shared between the developmental trajectories of root pericycle and procambium. However, there are still 19 genes unique to the ultimate development trajectory of xylem pole pericycle, including genes involved in cell differentiation (*AGP4/7/13/22/24*), cell cycle (*PATL1*), plant-type secondary-cell-wall biogenesis (*XTH15*), response to ABA (*CCH*), lignin biosynthesis (*AT1G55210* and *CAD4*), xylem differentiation (*XYP2*) and water transport (*PIP1C*) (**Figure 3H**). The result is consistent with previous findings that the xylem pole pericycle is an active cell dividing site (Dubrovsky et al., 2000).

In summary, the above results uncover previously uncharacterized developmental trajectories of plant tissues as tightly inner-connected gene expression programs that underlie the ontogeny of above- and belowground tissues.

### Orchestration of the cell fate specification by regulons

Gene expression is orchestrated by TFs in a cell-specific manner, and the cell fate specification can be characterized by unique expression patterns of gene regulatory networks (GRNs) where TFs and their target genes are generally co-expressed. Our integrated cell atlas allows the comparison of gene expression across the entire plant in a cell-type specific manner. To explore cell-type specific GRNs, SCENIC (Aibar et al., 2017) was used to predict active regulons based on co-expression analysis and TF motif enrichment. A regulon consists of a key TF regulator and its co-expressed target genes^28^. In total, we identified 628 regulons across all types of cells (**Figure 4A**, and **Supplemental Data Set S6**). As expected, the regulon activity of TFs and their expression intensity are highly correlated, confirming the reliability of regulon identification. We further used a published scATAC-seq dataset generated in root (Dorrity et al., 2021) to validate the identified regulons in root cells with a high accuracy (**Supplemental Figure S8**). Interestingly, the inferred regulons are generally cell-type specific (**Figure 4A**). For instance, the ANAC046 mediated regulon is specifically active in the root cap (**Figure 4B**). Consistent with this observation, ANAC046 was recently found to be a key regulator for cell death and for suberin biosynthesis in *Arabidopsis* root cap (Mahmood et al., 2019; Huysmans et al., 2018). The regulon containing JKD is highly active in the mature cortex. In line with its role in cortex regulation, JKD has been reported to interact with cell fate determinants SCR and SHR (Ogasawara et al., 2011) and to restrict *CYCIND6* expression in cortex, and mutation of *JKD* would lead to periclinal division in cortex(Welch et al., 2007). Similarly, the DOF5.6 and MYB3R4 regulons showed their explicit roles in the protophloem and proximal meristem cells, respectively. Besides, a number of regulons were shared among different cell types that are functionally correlated, as exemplified by the CRF10 and MYB40 regulons (**Figure 4B**). These two regulons showed high activity in the initiation cells and stele cells, respectively, indicating their crucial role in the development of specific cell types or in the transition between cell states.

**Figure 4:**
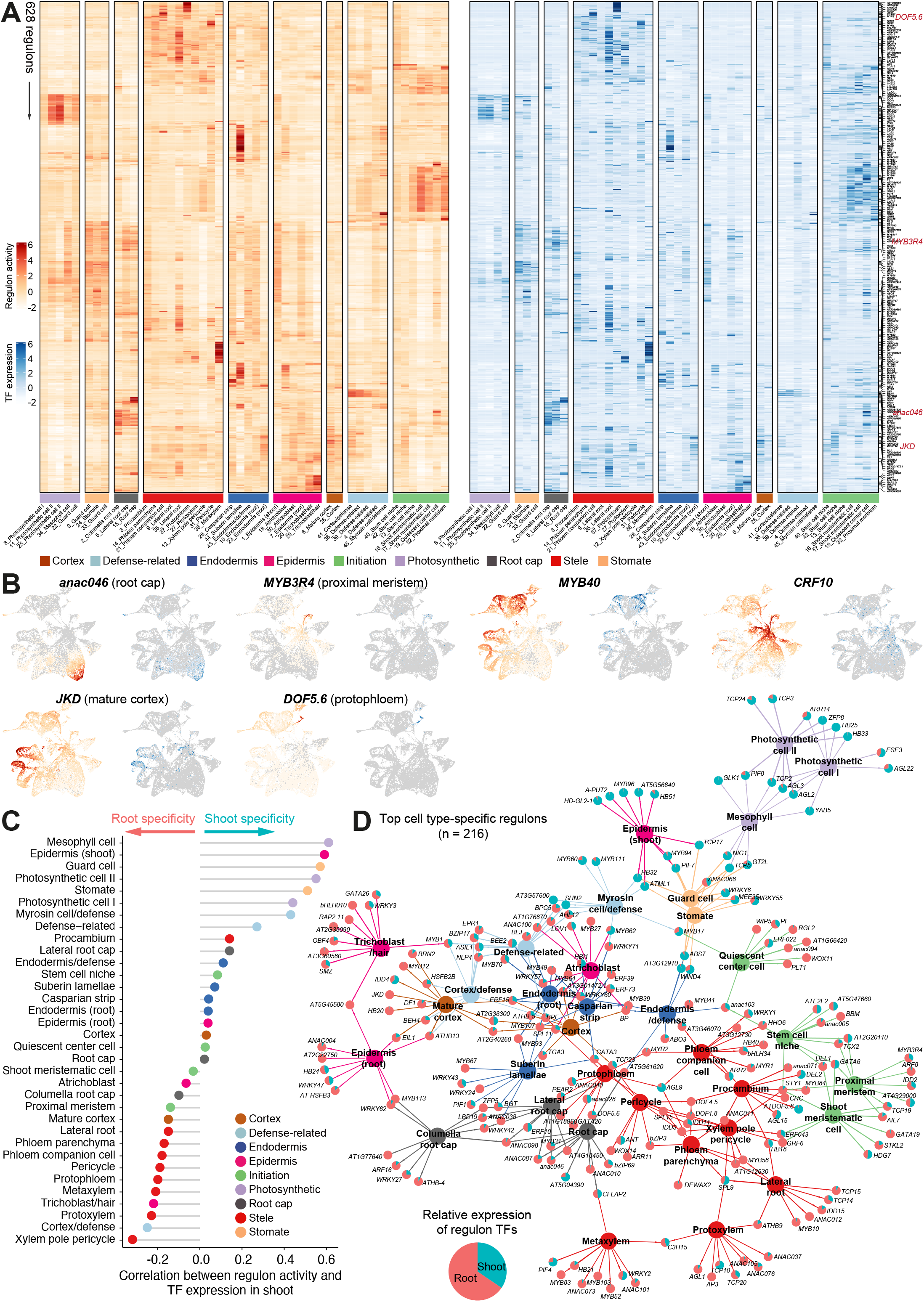
Inference of cell-type specific regulons by SCENIC. (**A**)Heatmaps showing the activity of 628 regulons per cell type (left), and the expression specificity of the corresponding regulon TFs (right). Top representative regulons in each cell types are highlighted. (**B**)UMAP plots depicting the activity and expression specificity of selected regulons. (**C**)Correlation of regulon activity and TF expression in shoot for each cell types. Positive correlations indicate cell types enriched in shoot and otherwise in root. (**D**)A network view of top ten representative regulons in each cell type, revealing specific and shared regulons across cell classes. Small nodes in pie charts represent the relative expression of regulon TFs in shoot and root. Large nodes indicate cell types which are colored according to the major cell classes.

To investigate the tissue-specific regulation of regulons, we performed correlation analysis between the regulon activity and the relative expression level of regulon TFs in either above- or belowground tissues for each cell type (**Supplemental Figure S9**). Intriguingly, the activity of regulons identified in the cell types in aboveground tissue, such as mesophyll cell, stomate cell and myrosin cells, showed positive correlations with the shoot expression of the corresponding shoot specific TFs, suggesting the shoot specificity of those regulons. In contrast, the activity of regulons from typical belowground tissue like lateral root and trichoblast is negatively correlated with the shoot expression pattern, indicating that they are root specific regulons (**Figure 4C**). Notably, the correlation for regulons from endodermis and initiation cells including quiescent center cells, shoot meristematic cells and root cap cells is almost neutral, indicating that these regulons are featured determinants for meristematic activity. As a proof of concept, we visualized the top ten active regulons in each cell type in a network view, where the relative expression of regulon TFs in root and shoot tissues were shown in pie charts (**Figure 4D**). In general, regulons were shared among cell types with similar functions or from the same major group. The identified regulons mainly expressed at the anatomy position of the plant (underground or aboveground) that correspond to the cell types they were assigned to. In QC center cells, PLT1 and WOX11 were highly activated; while in stem cell niche, BBM was one of the predominant regulators. All these three TFs can induce embryonic activity in plant cells (Kareem et al., 2015; Lowe et al., 2016; Liu et al., 2014). In short, the above analysis pinpoints key regulons underling plant cell fate specification.

### Dynamics of regulon-regulon associations

Since the dynamics of gene expression usually depend on cooperative binding of multiple TFs to the promoter regions (Jolma et al., 2015), we explored potential co-associations of regulons in different cell types. We adapted the Paired Motif Enrichment Tool (PMET)(Rich-Griffin et al., 2020) to detect pairs of TF binding motifs within the promoter regions of cell-type specific marker genes (Rich-Griffin et al., 2020). Only regulon TFs were used in the PMET analysis. The enriched TF pairs were subject to filtering using experimentally verified protein-protein interacted TFs, resulting in 1,295 reliable regulon-regulon pairs (**Figure 5A, Supplemental Data Set S7** and **8**). Taken the regulon pairs in photosynthetic cells as an example, TCP2, HY5, PTF1 and MEE35 (TCP4) were found to pair with a variety of other TFs to co-ordinately regulate photosynthesis related genes (Zheng et al., 2022; Toledo-Ortiz et al., 2014) (**Figure 5B**).

**Figure 5:**
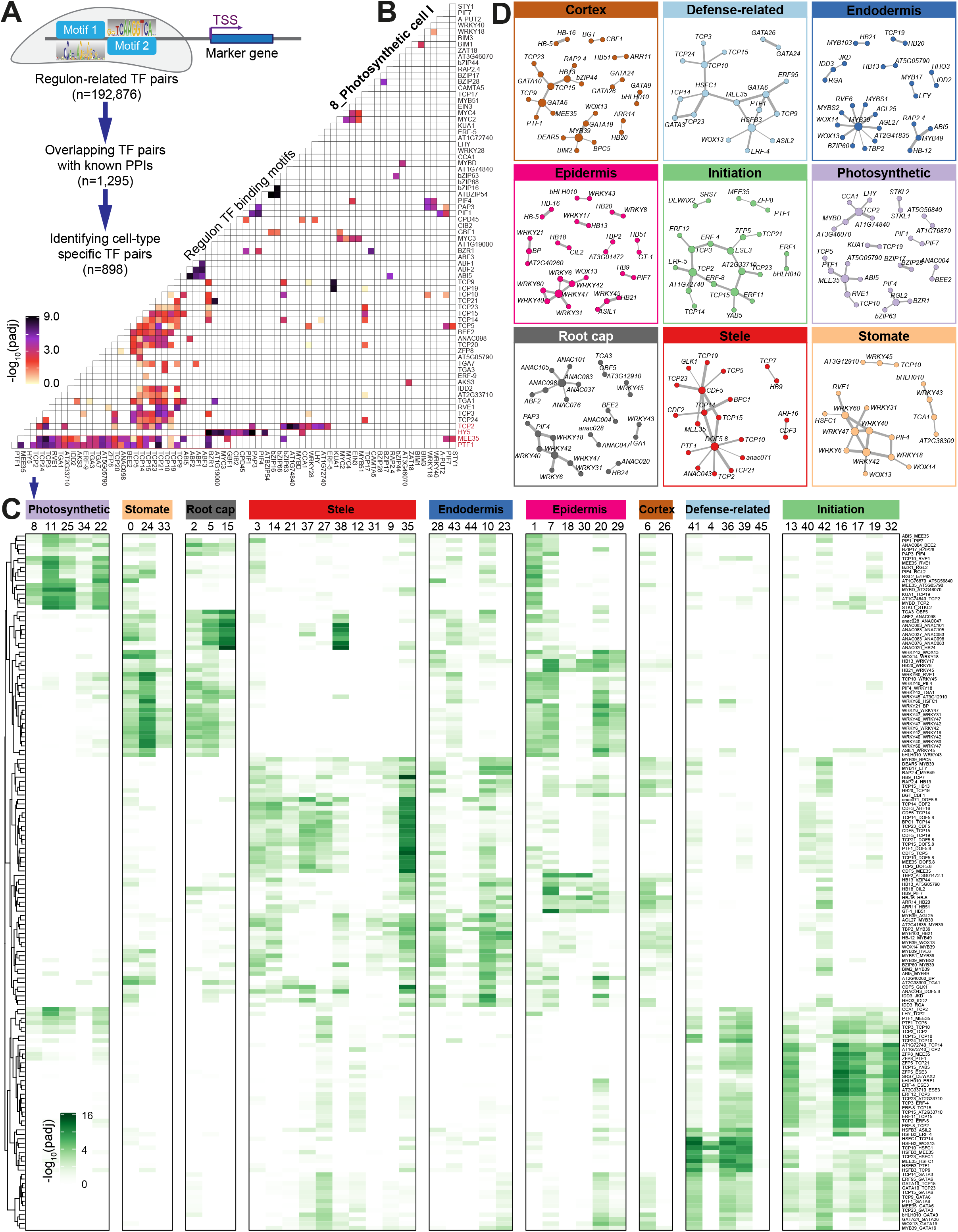
Combinational regulon-regulon pairs specifying cell type gene regulatory networks. (**A**)Workflow for identification of cell-type specific motif pairs. The diagram illustrates the paired-motif enrichment analysis in cell type identity genes, i.e., identifying pairs of TF binding motifs within the promoters of cell-type specific marker genes. (**B**)Heatmap representing an example of enrichment analysis of regulon pairs (in the photosynthetic cell cluster). (**C**)Heatmap showing the enrichment of representative regulon pairs (rows) whose enrichment scores are cell-type specific (columns). (**D**)Network view of representative regulon-regulon interactions for each major cell classes.

Interestingly, as the high cell-type specificity of active regulons (**Figure 4A**), most (69.3%, 898 out of 1,295) of the predicted co-associated regulons is also cell-type specific given that at least one of the interacting TFs was expressed cell-type specifically (**Figure 5A**). Indeed, we observed representative modules of regulon pairs which have clear cell-type-specific patterns (**Figure 5C-D** and **Supplemental Data Set S7**). Of note, regulon pairs in WRKY family were specifically enriched in stomatal cells, epidermis and root cap cells, in agreement with the biotic and abiotic stress related function of this TF family(Bakshi and Oelmüller, 2014). GATA-associated regulon pairs were absent in photosynthetic cells, while ERF-related regulon pairs showed high enrichment in initiation cells (**Supplemental Figure S10A-B**). Furthermore, we found that, among all the regulon pairs, the TCP family occupied more than 40% of pairs and presented in almost all cell types by co-association with other TFs (**Supplemental Figure S10C**), in line with the hub function of the TCP family members in various regulatory networks(Bemer et al., 2017). Our cell atlas allows looking into sophisticated TCP pairing patterns for gene regulation in different cell types. For instance, some of the TCPs, such as TCP3, TCP5 and TCP17, showed simple co-associations with TFs only in a few types of cells, while TCP2, TCP14, TCP15 and MEE35 (TCP4) tended to pair with multiple TFs in various cell types (**Supplemental Figure S10D**), suggesting diverse biological roles of the TCP family.

### Expression and function specificity of cell-type related gene regulatory networks

To examine how the cell-type specific regulons control dynamics of gene expression in different tissues, we constructed gene regulatory networks (GRNs) consisting of both the regulons and their target genes with tissue-specific gene expression (**Supplemental Data Set S9**). For visualization, only representative regulons in each cell types (as shown in **Figure 4D**) and their top five target genes with high specificity are displayed in the view of networks. The regulons were coloured according to their enriched major cell types and the target genes were shown in pie charts based on their relative expression levels in above- and underground tissues (**Figure 6A**). In general, there are similar expression patterns of target genes for the same regulon or regulons with identical cell-type specificity, indicating that the identified GRNs truly reflect the cell fate specification. A close-up view of the GRNs is shown in **Figure 6B**. The photosynthesis regulators AGL3, HB33 and HB25, which were reported to be involved in shoot morphogenesis (Hanano et al., 2020; Huang et al., 1995; Hong et al., 2011), were linked with target genes exclusively expressed in shoot. Interestingly, unlike other stele regulons, the target genes of the stele regulator ATDOF5.8 were highly expressed in aboveground tissues, including *WOX1* and *ER* which are well-known cell fate determinators in SAM development (Zhang et al., 2020). This suggests that ATDOF5.8 might play a key role in aboveground tissue development. IDD3 (also known as MAGPIE) was reported to interact with JKD, SCR and SHR and to regulate tissue boundaries formation in *Arabidopsis* root (Welch et al., 2007). Consistently, representative target genes of IDD3 were all root specified. Moreover, one of its target gene *AT1G62500* was recently identified as a marker gene of the meristematic cortex in root (Denyer et al., 2019).

**Figure 6:**
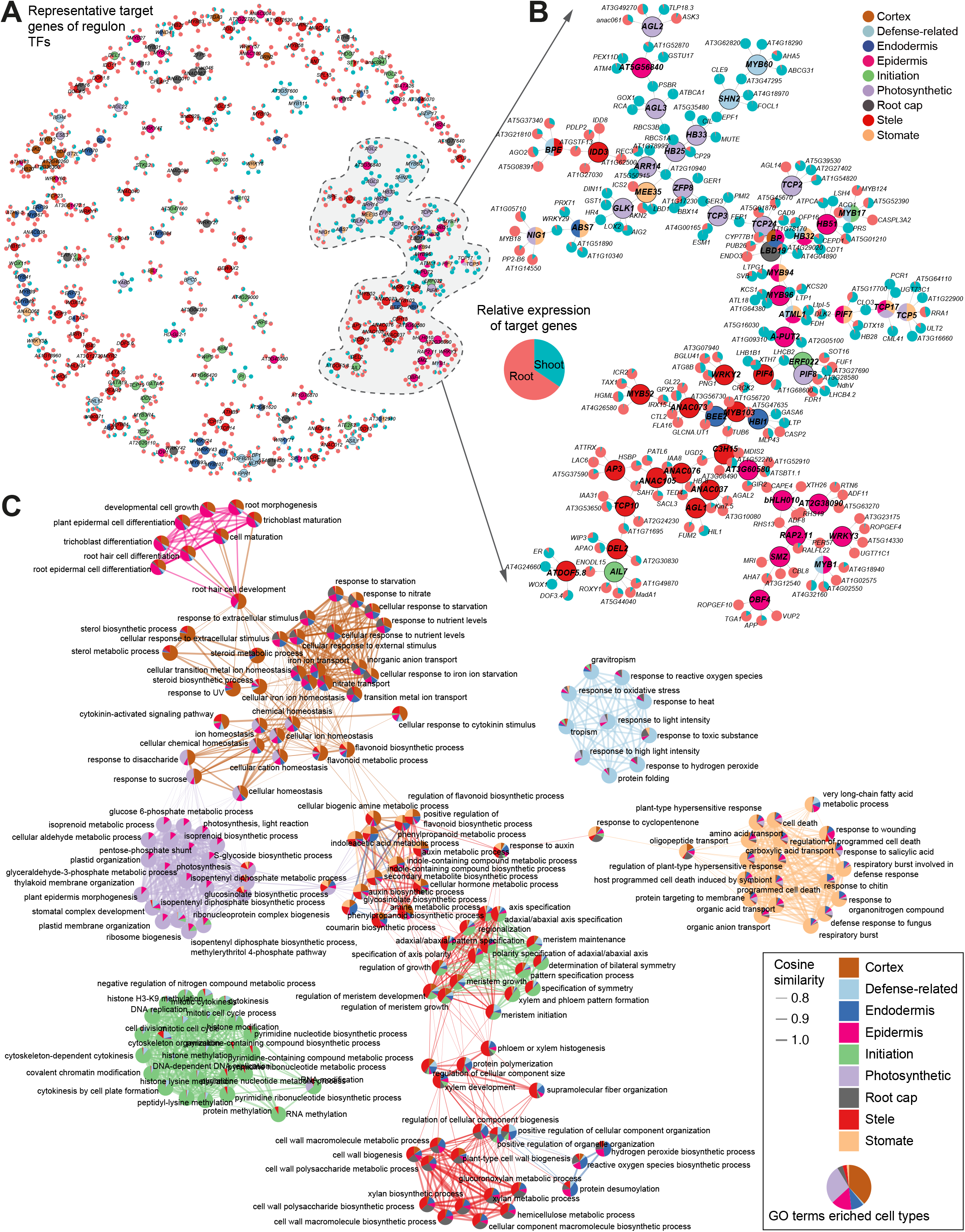
Expression and function specificity of cell type-related regulatory networks. (**A**)A network overview of representative target genes of cell-type specific regulons, revealing specific and common targets across regulons. Nodes with black circles represent TFs, colored according to the highly activated cell type; while nodes in pie charts indicate target genes. Note that only the top five target genes are shown for visualization purpose. (**B**)A detailed view of a subset of the network in (**A**). (**C**)A network analysis of GO terms (n=174; nodes) based on similarity of enrichment scores across cell types (edges), laid out using the spring-based algorithm by Kamada and Kawaial. GO terms are colored by the contributing cell types (pie chart by the fraction of enrichment scores in terms of −log_10_*P*).

Lastly, to study the functional specificity of cell-type GRNs, we performed GO enrichment analysis using all target genes for regulons identified in each cell type. We used GO term co-enrichment patterns among different cell types to cluster biological process terms. The resulting co-enrichment was shown in a network which synoptically revealed several densely connected communities of biological processes with meaningful similarities (**Figure 6C**). Unsurprisingly, environmental stimuli related biological pathways such as response to light intensity, gravitropism and oxidative were specially enriched in defense-related cells, supporting a hypothesis that defense-related cells might be a previously uncharacterized cell type in response to stimulation of external environment. Similarly, biological processes related to meristem activity and epigenetic regulation were highly enriched in initiation cells, consistent with their biological roles in cell fate reprogramming. Notably, some cell types shared a common set of GO terms with biologically meaningful similarities. For instance, nutrient level response related process were enriched in cortex, endodermis, epidermis and stele cells, consistent with the knowledge that absorption and transportation of nutrients are through concentric layers of those tissues (Barberon, 2017).

## Discussion

Recently, empowered by the burst of single-cell sequencing techniques, references of single cell atlas across tissues or of the whole organisms have largely been generated in various model species especially in mammalian species (Qu et al., 2022; Han et al., 2020, 2018; Jones et al., 2022). To this end, the Human Cell Atlas (HCA) project aims to provide comprehensive reference maps of all human cells (Regev et al., 2017; Rozenblatt-Rosen et al., 2017). In the plant field, similar attempt such as the Plant Cell Atlas (PCA) framework has been proposed (Jha et al., 2021; Rhee et al., 2019). In fact, dozens of studies in plant sciences have fully embraced the single cell sequencing techniques, leading to generation of significant amounts of single-cell transcriptome datasets in specific tissues or organs over the past years (Seyfferth et al., 2021; Shaw et al., 2021). These advances have not only expanded our knowledge about plant cell compositions, but also laid the groundwork for PCA construction. One of the major challenges for PCA is to generate a reference cell map that includes all the cell types (or as much as possible) in various tissues (Cuperus, 2022). However, current plant cell atlases are constructed in specific tissues and we still lack of a reference cell map of the entire plant. Therefore, collecting cells generated in representative tissues from different labs and development of data integration analysis pipelines would be the most effective way to achieve an “universal” annotation of cell types in plants. Consequently, here we have piloted an integration analysis of scRNA-seq data in *Arabidopsis* seedlings which represents a reference atlas at the early plant development stage. Our study not only provides valuable resources for PCA exploration, but also presents a guideline for integrative analysis of plant single cell datasets.

The power of integrative analysis of scRNA-seq data has been proved by the discovery of novel cell types (**Figures 1C** and **2A-B**). Defense-related cells are such a novel but rare cell type in our analysis. Although plant immune systems lack of mobile immune cells, a group of cells that have specific response towards external stimulus have brought up to our particular attention. We hypothesise that transcription feature of these rare cell types can easily be diluted in bulk RNA-seq analysis, and are not detectable when the sample size is not large enough to extract signal from noise. Therefore, the defense-related cells can only be distinguished in our large-scale data integration then enough rare cells are enriched into a specific population for cell type detection. Nevertheless, experimental validations based on RNA-fluorescence in situ hybridization (FISH) and marker genes fused with reporters need to be conducted to further confirm the findings of novel cell types and marker genes.

The development of plant organs and tissues relies on the accurate regulation of cellular differentiation. Cell fate specification is a progress of differential gene expression typically mediated by multiple TFs in a coordinate way (Drapek et al., 2017; Reiter et al., 2017). The comprehensive cell map of *Arabidopsis* presented in the current study enables to delineate the architecture of GRNs underlying plant cell fate decisions at an unprecedented resolution (**Figures 4D** and **5**). The resulting cell-type specific GRNs will help to generate hypotheses to address specific mechanisms of differentiation. Our results also imply some novel function of previously identified TFs. For instance, *ANAC020* was known as a regulator in the differentiation of phloem (D et al., 2015). Our analysis showed that *ANAC020* specifically paired with *MYB1* and *BIM1* in metaxylem and protopholem cells (**Figure 5C** and **Supplemental Data Set S8**). Interestingly, *MYB1* and *BIM1* are both key regulators of xylem and phloem cell differentiation from vascular stem cells (Y et al., 2005; Saito et al.). This pairing pattern indicates the hidden role of *ANAC020* through coordination with other cell identity TFs to determine the fate of vascular cells.

Big dreams start small. Although references of plant cell atlases are far from complete, the integrative cell atlas constructed by our study provides a first reference cell map at the level of the entire plant. The generated cell-type specific GRNs in *Arabidopsis* will lay the groundwork for comparative genomics of cell fate processes in other plants.

## Methods

### scRNA-seq data collection

scRNA-seq datasets (n = 45; **Supplemental Data Set S1**) generated in *Arabidopsis juvenile* seedlings by 10X Genomics or Drop-seq technologies were collected from previous studies (Liu et al., 2020; Lopez-Anido et al., 2021; Denyer et al., 2019; Wendrich et al., 2020; Zhang et al., 2019; Long et al., 2021; Jean-Baptiste et al., 2019; Ryu et al., 2019; Shulse et al., 2019; Farmer et al., 2021). We reanalysed these scRNA-seq data using the raw sequencing data in the fastq format, which were downloaded from the Sequence Read Archive (SRA) database (https://www.ncbi.nlm.nih.gov/sra/).

### Pre-processing of raw sequencing data

The raw 10X Genomics and Drop-seq fastq files were aligned, filtered and counted using the Cell Ranger pipeline version 3.1.0 (10x Genomics) and the Drop-seq tool version 1.13 (https://github.com/broadinstitute/Drop-seq) respectively. The *Arabidopsis* reference genome (version TAIR10) and the corresponding GTF annotation file (version Araport11) were downloaded from the *Arabidopsis* Information Resource (TAIR) database (https://www.arabidopsis.org/), followed by genome index built and filtered reads alignment using STAR (version 2.7.3a), which used as an aligner in Cell Ranger and Drop-seq tools. Specifically, for the alignment of reads generated from single nucleus, in order to accommodate the expression of precursor RNAs that contain introns, the intron regions of each read were removed and realigned to the reference genome. After barcodes and UMIs counting, the feature-barcode matrixes generated from Cell Ranger and the digital expression matrixes returned by Drop-seq tools, all of which with each unique molecular identifiers (UMIs) for every detected gene as a row and per valid cell barcode as a column, were used for subsequent analyses.

### Integration of single cell data

Feature-barcode count matrices for each sample were processed with *Seurat* package (v4.0.0) (Hao et al., 2021). Two types of cells were removed from the analysis: the ones expressed less than 200 genes or the ones have more than 10% of mitochondrial gene expression in UMI counts. “vst” in “FindVariableFeatures” function were utilized to determine the Top 3000 most variably expressed genes, “mt.percent” in“ScaleData” was further used with regression on the proportion of mitochondrial UMIs. “Two-stepwise strategy” was applied for scRNA-seq data integration. Firstly, scRNA-seq datasets from a specific study were aligned with canonical correlation analysis (CCA), then Robust Principal Component Analysis (RPCA) method in Seurat was used for integration of scRNA-seq data from different studies.

### Cell clustering and annotation

“RunPCA” function was used to compute the top 30 principal components (PCs) using top variably expressed genes on visualization purposes. Clustering was performed with integrated expression values based on shared-nearest-neighbor (SNN) graph clustering (Louvain community detection-based method) using “FindClusters” with a resolution of 0.8. Cell clusters were visualized using UMAP (Uniform Manifold Approximation and Projection).

Differential expressed genes (DEGs) in each cluster were identified using “FindAllMarkers” with non-parametric Wilcoxon rank sum test method. For cell annotation, we firstly constructed a cell type-specific marker genes database via collected annotated marker genes from previous published dataset, including single-cell studies as well as abundant gene expression studies (**Supplemental Data Set S3**). Then, the defined cell types of our detected conserved cluster-specific marker genes were retrieved from the cell type-specific marker genes database. Finally, we consulted the documented function and verified the expression pattern of each cell-specific marker gene. The expression pattern of cell known and expected cell type-specific marker genes across clusters were visualized to further confirm the cell type of each cluster.

### Cell type deconvolution

We first generated a cell-type expression matrix of top 100 marker genes from scRNA-seq data. Using this gene expression matrix, we then applied CIBERSORTx (Newman et al., 2019) with default parameters to deconvolute cell-type abundance from 96 bulk mRNA-seq datasets collected from the PPRD database (Yu et al., 2022) (**Supplemental Data Set S5**).

### Single cell development trajectory reconstruction

The Monocle (v.2.8.0) package based on R (v4.0.0) was used to infer the process of cell differentiation and cell fate determination. The DEGs identified by Seurat were used to sort the cells in pseudotime order. Then, we performed “DDRTree” to reduce the dimension into two components and using “plot_cell_trajectory” to visualize the cell trajectory. For priori “beginning” specification of the trajectory tree, we run “orderCells” again with a specific cell cluster set. Finally, BEAM was applied to analyse the DEGS depending on each branch or pseudotime, the genes that significantly branch-dependent were plotted via the function of “plot_genes_branched_heatmap”.

### Gene regulatory network inference and regulons discovery

TF DNA binding motifs of *Arabidopsis* were downloaded from the JASPAR(Fornes et al., 2020) and CIS-BP (Weirauch et al., 2014) databases and subjected to redundancy filtering. The cisTarget database was constructed according to the SCENIC (Aibar et al., 2017) protocol (https://github.com/aertslab/create_cisTarget_databases). We used pySCENIC 0.11.2 (https://github.com/aertslab/pySCENIC) (Aibar et al., 2017) to infer gene regulatory network. Briefly, SCENIC contains three steps: (1) identify co-expression modules between TF and the potential target genes; (2) for each co-expression module, infer direct target genes based on motif enrichment of the corresponding TF. Each regulon is then defined as a TF and its direct target genes; (3) calculate the Regulon Activity Score (RAS) in each single cell via the area under the recovery curve.

### Regulon-regulon co-association analysis

We used Paired Motif Enrichment Tool (PMET; https://github.com/kate-wa/PMET-software)(Rich-Griffin et al., 2020) to detect the co-localisation of pairs of TF binding motifs within the promoters of our cell identity and cell type-specific differentially expressed genes. Only regulon TFs were included in the analysis. Importantly, experimentally verified protein-protein interaction (PPI) information of *Arabidopsis* from the databases of BIND (http://bind.ca), BIOGRID (https://thebiogrid.org/), IntAct (https://www.ebi.ac.uk/intact/home) and MINT (https://mint.bio.uniroma2.it/) were collected and utilized to filter the predicted regulon-regulon TF pairs.

### Gene set enrichment analysis

Gene ontology (GO) enrichment analysis for a specific gene set was performed using the *clusterProfiler*(Wu et al., 2021) package.

### Topic modeling of cell-type specific gene expression programs

In order to identify cell-type specific gene expression programs (GEPs) with biological meanings, we used the *CellFunTopic* framework to infer enriched biological pathways for DEGs among different cell types. Specifically, *CellFunTopic* scores the activity of each functional gene set defined by GO vocabularies using the gene set-scoring method of Gene Set Enrichment Analysis (GSEA) (Subramanian et al., 2005). The resulting enrichment scores in terms of clusters-by-gene-set scoring matrix is subjected to topic modeling. *CellFunTopic* applies latent Dirichlet allocation (LDA) (Blei et al., 2003) with variational expectation maximization (VEM) (Nasios and Bors, 2006) to factorize the scoring matrix into a clusters-by-topics matrix *θ* (the probability of a cluster belonging to a topic) and a topics-by-gene-set matrix *ϕ* (the contribution of a topic within a cell cluster). The results are highly interpretable: the top topics under each cell cluster directly reveal the specificity of biological programs belonging to a particular cell type, which is analogous cell-type GEPs.

### Statistical analysis

If not specified, all statistical analyses and data visualization were done in R (version 4.0.0). R packages such as *ggplot2* and *igraph* were used for graphics.

## Data availability

The processed and integrated single cell data in this study can be retrieved and viewed at https://biobigdata.nju.edu.cn/plantScGRN/.

## Code availability

CellFunTopic is available at https://github.com/compbioNJU/CellFunTopic. R codes used to analyze data and generate figures are available upon reasonable request to the corresponding authors.

## Acknowledgements

This work is supported by the National Natural Science Foundation of China (No. 32070656), the Nanjing University Deng Feng Scholars Program and the Priority Academic Program Development (PAPD) of Jiangsu Higher Education Institutions and China Postdoctoral Science Foundation funded project (No.2022M711563).

## Author contributions

D.C. conceived the study. D.C, W.Y. and C.Z. supervised the study. S.C. and D.C. performed data analyses with supported from C.H., X.Z., R.Y., Y.L. and W.F.. D.C., X.Z. and C.H. wrote the manuscript with input from W.Y. and C.Z.. All authors reviewed and approved the submitted version.

